# Intercellular trafficking of latent TGF-β1 converts integrin-expressing cells into signaling relays

**DOI:** 10.64898/2026.11.09.765446

**Authors:** Aravinda-Bharathi Ramakrishnan, Xiang Li, Jude Kim, Viet Q. Le, Timothy A. Springer

## Abstract

TGF-β1 is secreted as a latent complex tethered to the cell surface by milieu molecules such as GARP, and its activation requires direct engagement by integrin-expressing cells. How this interaction converts the latent complex into signaling-competent growth factor (GF) remains unclear. Here we show that contact between GARP/TGF-β1-presenting cells and cells expressing integrin αVβ6 or αVβ8 triggers rapid trogocytosis of the latent complex, both in transfected cells and two cancer cell lines that natively express these integrins. Independent labeling of the prodomain, GF, and GARP shows that all three components are co-transferred to the integrin-expressing cell, and a TGF-β1 mutant (RGE) that cannot bind integrins shows little transfer. Using a reporter system that tracks pathway activation in presenting, integrin-expressing, and bystander cells, we find that GF-neutralizing antibodies block downstream signaling more effectively in bystander cells than in the first two cell types that participate in trogocytosis. Blocking cleavage between the prodomain and the GF blocks signaling in bystander cells while still permitting signaling in trogocytosing cell types. Finally, we integrin-expressing cells that have taken up latent TGF-β1 by trogocytosis can activate it and signal to bystander cells after the TGF-β1 cells are removed, indicating that trogocytosis converts integrin-expressing cells into TGF-β signaling relays. These findings identify a role for trogocytosis in regulating the specificity of TGF-β1 activation and signaling, and reveal a novel function of integrins in controlling protein localization.

## Introduction

Transforming growth factor-β1 (TGF-β1) is a signaling protein that plays a vital role in immune homeostasis, in addition to regulating the proliferation and differentiation of many cell types ^1,2^. Mice lacking *Tgfb1* function develop lethal multiorgan inflammation after birth^3,4^. These mice also exhibit many defects of vasculature ^5^. In addition to its immunoregulatory and developmental roles, TGF-β1 is a major driver of fibrotic diseases and immune resistance in cancer ^2,6,7^.

TGF-β1 is produced as a latent procomplex in which a “prodomain” (also known as the Latency Associated Peptide/LAP) is non-covalently associated with the “growth factor” (GF) domain. Henceforth we use “TGF-β1” to refer to the full-length TGF-β1 protein comprising of a prodomain in complex with the GF. The prodomain prevents the GF from binding to TGF-β receptors to trigger downstream signaling ^8,9^. TGF-β1 can be activated through interactions with integrins αVβ6 and αVβ8. The prodomain of TGF-β1 contains an RGD motif that is bound by both integrins ^10,11^. Mice bearing mutations of the RGD to block integrin binding ^12^, or mutation and inhibition of integrins αVβ6 and αVβ8^13^ phenocopy *Tgfb1* null mice, establishing the essential role of integrin-mediated activation in TGF-β1 function.

TGF-β1 is expressed in complex with “milieu anchor” proteins through inter-protein disulfide linkage ^8^. The milieu anchors GARP (LRRC32) and LRRC33 (NRROS) are transmembrane proteins that display TGF-β1 on the surfaces of immune cells. Regulatory T cells (Tregs) and platelets express GARP/TGF-β1 complexes^14–16^, while macrophages, microglia, and other myeloid-derived lineages express LRRC33/TGF-β1 complexes ^17^. GARP/TGF-β1 presented by Tregs and LRRC33/TGF-β1 presented by tumour-infiltrating myeloid cells both serve as sources of TGF-β1 in cancers^18,19^. This is thought to be activated by integrin αVβ8 expressed by tumor cells and immune cells such as Tregs and dendritic cells ^20–23^.

Although molecular, biochemical, and genetic lines of evidence converge on integrin binding to the prodomain being essential for activation, where this activation occurs is still an open question. Although the effects of loss of TGF-β1 have been studied in different cell types, how downstream pathway activation occurs in different cells in unknown. What is the fate of the prodomain and milieu anchor after the TGF-β1 molecule is activated and the GF released? Does the GF simply diffuse away from the site of activation to target cells? We tracked the members of the GARP/TGF-β1 complex and found they were transferred intact to cells expressing activating integrins by direct prodomain-integrin interactions through the RGD motif. This transfer, through a process called “trogoyctosis”, allows the activation of downstream signaling in TGF-β1 producing cells and integrin-expressing TGF-β1 activating cells. It does not require the cytoplasmic tails of integrins αVβ6 and αVβ8 or any of their known cytoplasmic adaptor proteins. Trogocytosis of TGF-β1 results in cells that are TGF-β1 producers or activators being less sensitive to TGF-β1 GF blocking antibodies than cells that are bystanders. We show that cancer cells can act as “signal relays” for TGF-β1, taking up latent complexes when they encounter TGF-β1-expressing cells, and subsequently releasing active GF to nearby cells even in the absence of the cell that originally produced TGF-β1. These results reveal a new mechanism for restricting and relaying TGF-β1 activation and suggest how immunosuppressive tumor microenvironments may be established.

## Results

### Integrin-dependent intercellular transfer of TGF-β1 complex components

Cell-culture models in which integrin-and TGF-β-expressing cells are co-cultured have been used to study the mechanistics of TGF-β activation ^14,24^. To understand the fate of different components of the latent GARP/TGF-β1 complex during TGF-β activation by integrins, we developed a system to fluorescently label and track the localization of each component of the complex. We tracked the prodomain using an N-terminal Flag tag to monitor the expression of TGF-β1 and TGF-β2^24^. For labelling the TGF-β1 growth factor (GF), we utilized amber codon suppression^25–27^ to site-specifically incorporate the non-canonical amino acid trans-Cyclooct-2-en-L-Lysine (TCO*-lys) in place of lysine 344 (K344) in the GF domain. K344 was selected for TCO*-lys incorporation because it is not involved in interactions with GARP, the prodomain, or TGF-β receptors based on structural analysis ^28,29^;^30^. TCO*-lys incorporated at residue 344 was site-specifically labeled with the tetrazine-functionalized fluorophore LD655^31,32^. Co-culture of cells expressing LD655-labelled GARP/TGF-β1 with integrin αVβ6 and αVβ8-expressing cells produced greater TGF-β activation than mock transfected cells in a reporter assay^18,24^ confirming functionality of the K344* mutant construct (Supplementary fig. 1A). To label GARP, we fused a Halotag^33^ to its N-terminus, immediately following the signal peptide. A reporter assay confirmed that the Halotag did not interfere with GARP/TGF-β1 activation (Supplementary fig. 1B).

To distinguish between cells transfected with GARP/TGF-β1 and those transfected with integrins after co-culture, we labelled integrin transfectants with the cytoplasmic dye CellTracker Green CMFDA prior to co-culture. This allowed us to distinguish integrin-expressing cells from GARP/TGF-β1 cells in co-cultures by flow cytometry on the basis of CMFDA signal (Fig. 1A, Supplementary fig. 1C-E). We first used this system to track the movement of the GF. LD655-labeled GARP/TGF-β1 K344*-expressing cells were co-cultured for 10 minutes with CMFDA-labeled Expi293F cells that were mock-transfected or transfected with integrin αVβ6, αVβ8, or αVβ1. Co-cultures were then placed on ice and fluorescence was measured by flow cytometry. In overnight reporter assays, αVβ6 and αVβ8 transfectants, but not αVβ1 transfectants, activate TGF-β1^14,34^. Consistent with this, LD655 signal was lost from GARP/TGF-β1 cells co-cultured with αVβ6 or αVβ8 transfectants, but not from cells co-cultured with mock or αVβ1 transfectants (Fig. 1B-D). This was accompanied by an increase in LD655 signal in αVβ6 and αVβ8 transfectants compared with mock and αVβ1 transfectants, suggesting that significant GF is transferred from TGF-β1-expressing cells to integrin-expressing cells within 10 minutes of contact.

**Figure 1.**
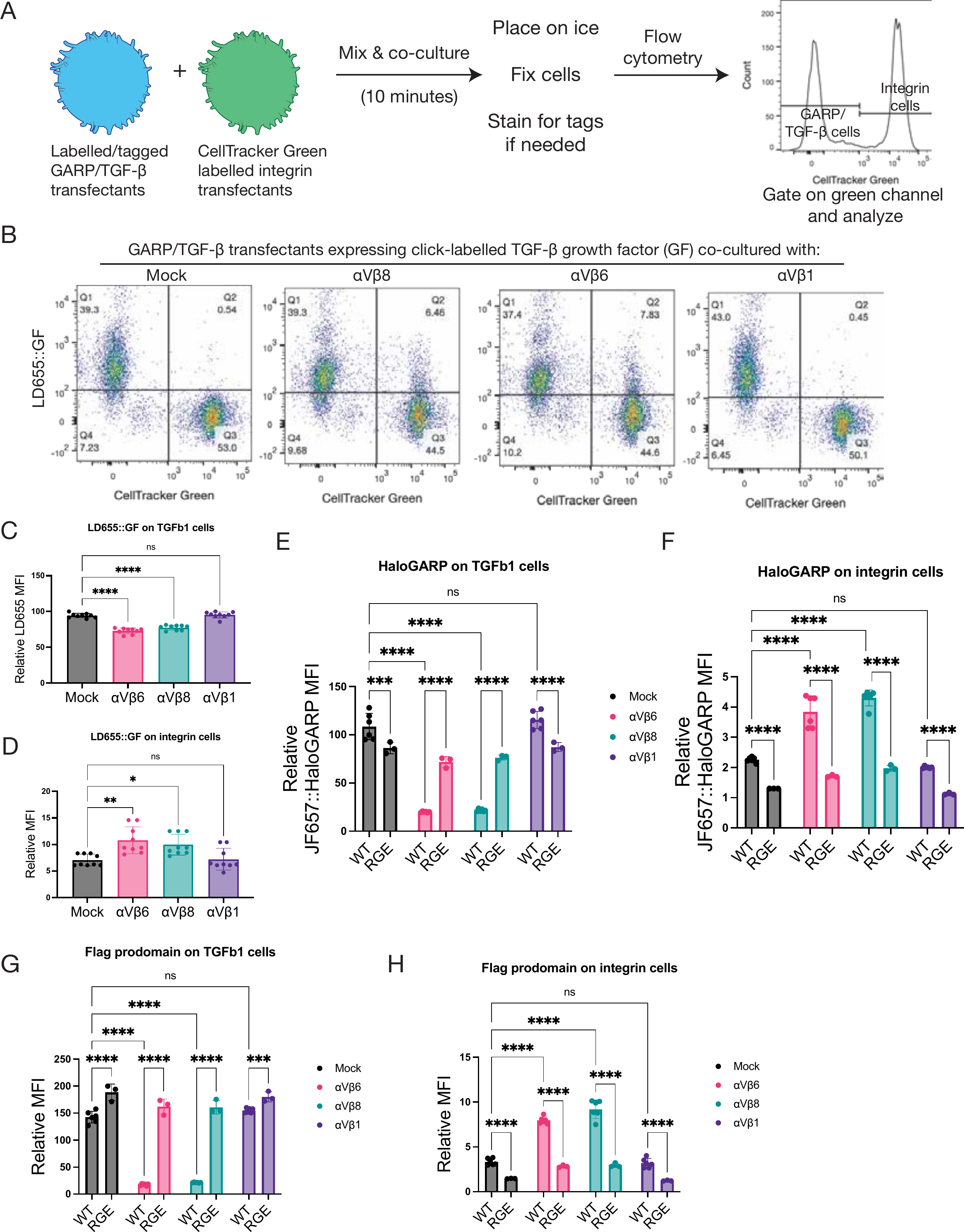
Interaction between GARP/TGF-β1-expressing cells and cells expressing TGF-β1-activating integrins results in the transfer of GARP, TGF-β1 prodomain, and growth factor (GF) to integrin-expressing cells. A. Schematic of flow cytometry-based trafficking assay to study localization of GARP/TGF-β1 complex members during TGF-β activation. B. Representative dot plot showing CellTracker Green CMFDA and LD655 signals from co-cultures of LD655-labelled GARP/TGF-β1 cells with CellTracker-labelled Expi293F cells transfected with indicated expression constructs. C. Quantification of LD655::GF signal in GARP/TGF-β1 cells co-cultured with indicated transfectants showing depletion by integrins αVβ6 and αVβ8. D. LD655::GF signal in integrin cells co-cultured with GARP/TGF-β1 cells showing transfer of GF to αVβ6 and αVβ8-expressing cells. E. Quantification of JF657::Halo-GARP signal in cells transfected with Halo-GARP and either TGF-β1 wild-type or RGE and incubated with indicated integrin transfectants. F. JF657::Halo-GARP signal in integrin transfectants co-cultured with Halo-GARP/TGF-β1 WT or RGE. Halo-GARP is transferred only to αVβ6 and αVβ8-expressing cells and transfer is blocked by the RGE mutation. G, H. Quantification of Flag-prodomain signal in GARP/TGF-β1 (G) and integrin-expressing cells (H) as in E and F, showing transfer of prodomain to αVβ6 and αVβ8-expressing cells.

We similarly addressed what happens to the prodomain and GARP in the same assay. Halo-GARP on Halo-GARP/Flag-TGF-β1 transfectants was fluorescently labeled with the JF657 HaloTag ligand^35^. After 10 min of co-culture with mock or integrin transfectants, the Flag-prodomain was labeled by immunostaining on ice. Because immunostaining was done without fixation or permeabilization, only surface-exposed Flag-prodomain was labeled. As a negative control, we used a TGF-β1 construct with an Asp-to-Glu mutation in the RGD integrin-recognition motif of the prodomain (RGE mutant), which disrupts binding of integrins αVβ6 and αVβ8 and subsequent activation^34,36,37^. Halo-GARP was strongly depleted from GARP/TGF-β1 transfectants co-cultured with integrin αVβ6 or αVβ8 transfectants, but not with mock or αVβ1 transfectants (Fig. 1E). This was accompanied by the transfer of GARP into integrin αVβ6 and αVβ8 transfectants (Fig. 1F). This depletion and transfer were strongly attenuated with the RGE mutant (Fig. 1E,F), indicating that direct binding of integrins to the TGF-β1 prodomain drives this process. Tracking the prodomain with an antibody gave similar results (Fig. 1G,H).

Thus, while the loss of the TGF-β1 GF from GARP/TGF-β transfectants and transfer to cells expressing integrin αVβ6 and αVβ8 could have resulted from GF release from the pro-complex, parallel loss and transfer to cells expressing αVβ6 and αVβ8 of the pro-complex suggest a distinct and previously undescribed process of transfer of intact TGF-β1 pro-complexes to integrin bearing cells. Furthermore, our results show this process is highly specific, as it requires an intact RGD motif on TGF-β1 and occurs with integrins αVβ6 and αVβ8 and not αVβ1.

We next sought to rule out overexpression artifacts and to test whether endogenous levels of integrin expression were sufficient to mediate the transfer of GARP/TGF-β1 complexes into integrin-expressing cells. We examined two cancer cell lines: LN229, a glioblastoma cell line that expresses integrin αVβ8^38^, and BT-20, a breast cancer cell line that natively expresses integrin αVβ6. We confirmed integrin expression by immunostaining with function-blocking antibodies specific to αVβ8 and αVβ6 and a function-blocking pan-αV-targeting antibody (Supplementary fig. 2). Similar to integrin transfectants, we observed the transfer of GARP and Flag-prodomain to LN229 and BT-20 cells, coincident with their depletion from GARP/TGF-β1 transfectants (Fig. 2A-H). Transfer of GARP and Flag-prodomain to LN229 and BT-20 cells was blocked by appropriate integrin-specific antibodies as well as the pan-αV-neutralizing antibody. These results show that GARP/TGF-β1 complex components are transferred to integrin-expressing cells in a manner dependent on TGF-β1-activating integrins.

**Figure 2.**
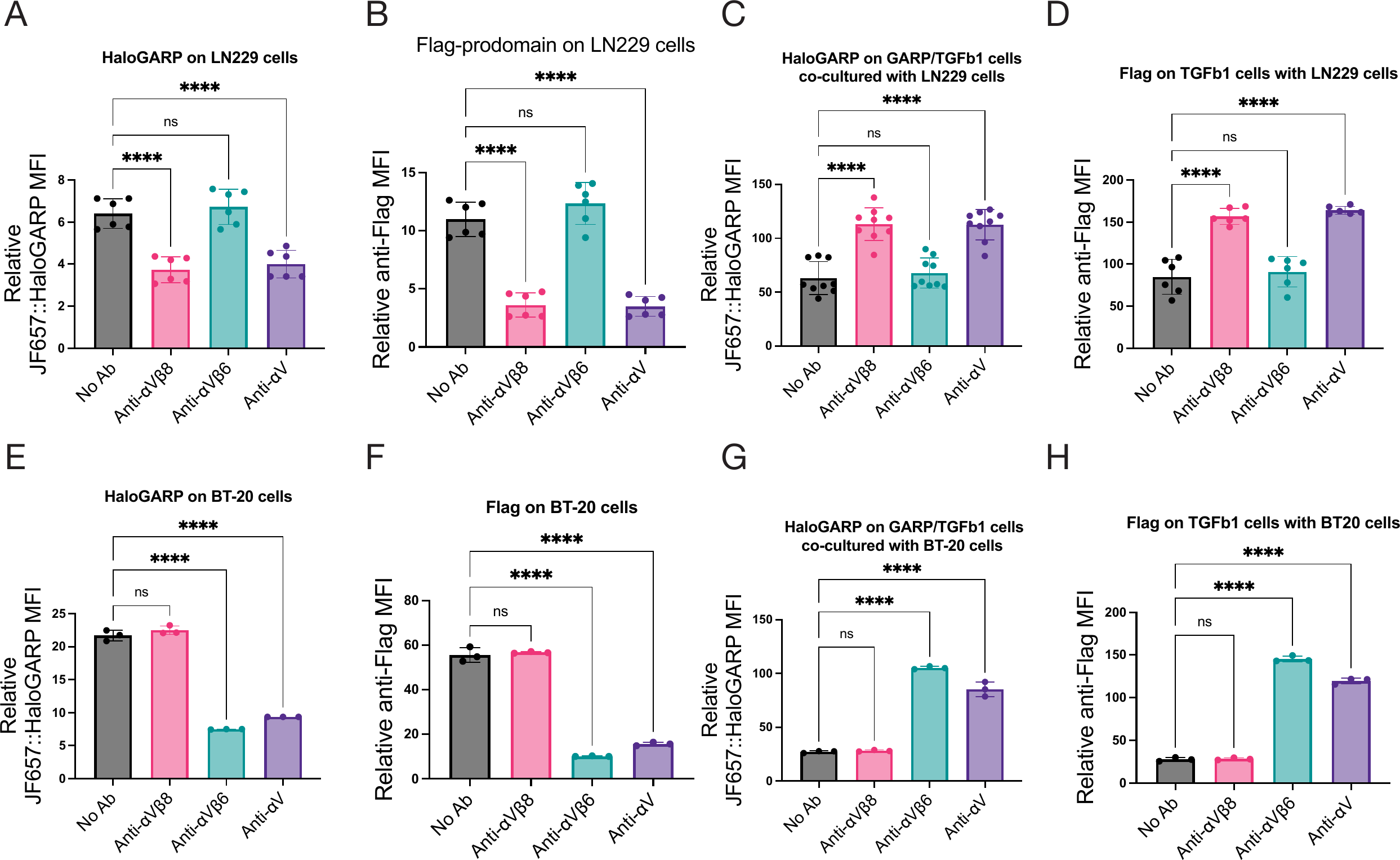
Cancer cells that natively express integrins αVβ6 or αVβ8 take up GARP/TGF-β1 complexes from cells that express them. A,B. Quantification of Halo-GARP and prodomain signals in αVβ8-expressing LN229 cells co-cultured with GARP/TGF-β1 transfectants showing that transfer is blocked by αV and αVβ8-function blocking antibodies. C,D. Quantification of Halo-GARP and prodomain signals in GARP/TGF-β1 transfectants co-cultured with LN229 cells. E,F. Quantification of Halo-GARP and prodomain signals in αVβ6-expressing BT-20 cells co-cultured with GARP/TGF-β1 transfectants showing that transfer is blocked by αV and αVβ6-function blocking antibodies. G,H. Quantification of Halo-GARP and prodomain signals in GARP/TGF-β1 transfectants co-cultured with BT-20 cells. All MFIs are expressed relative to the staining on labeled GARP/TGF-β1 transfectants without co-culture.

### Intact TGF-β1 complexes are transferred to integrin-expressing cells

Having found that GARP, prodomain, and GF are all trafficked to activating integrin-expressing cells, we asked whether the complex was transferred intact. We also asked whether the material was internalized or remained on the surface of integrin-expressing cells. To address both questions, we co-cultured GARP/TGF-β1 cells with CMFDA-labelled mock, αVβ6 and αVβ8 transfectants, and stained them with the TW7-28G11 antibody, which recognizes only the intact latent TGF-β1 complex and not the prodomain alone^39^. Integrin-expressing cells were positive for latent TGF-β, while mock cells were not (Fig. 3A,D). Consistent with their transfer to integrin-expressing cells, latent TGF-β1 complexes were depleted from surfaces of GARP/TGF-β1 transfectants (Fig. 3E).

**Figure 3.**
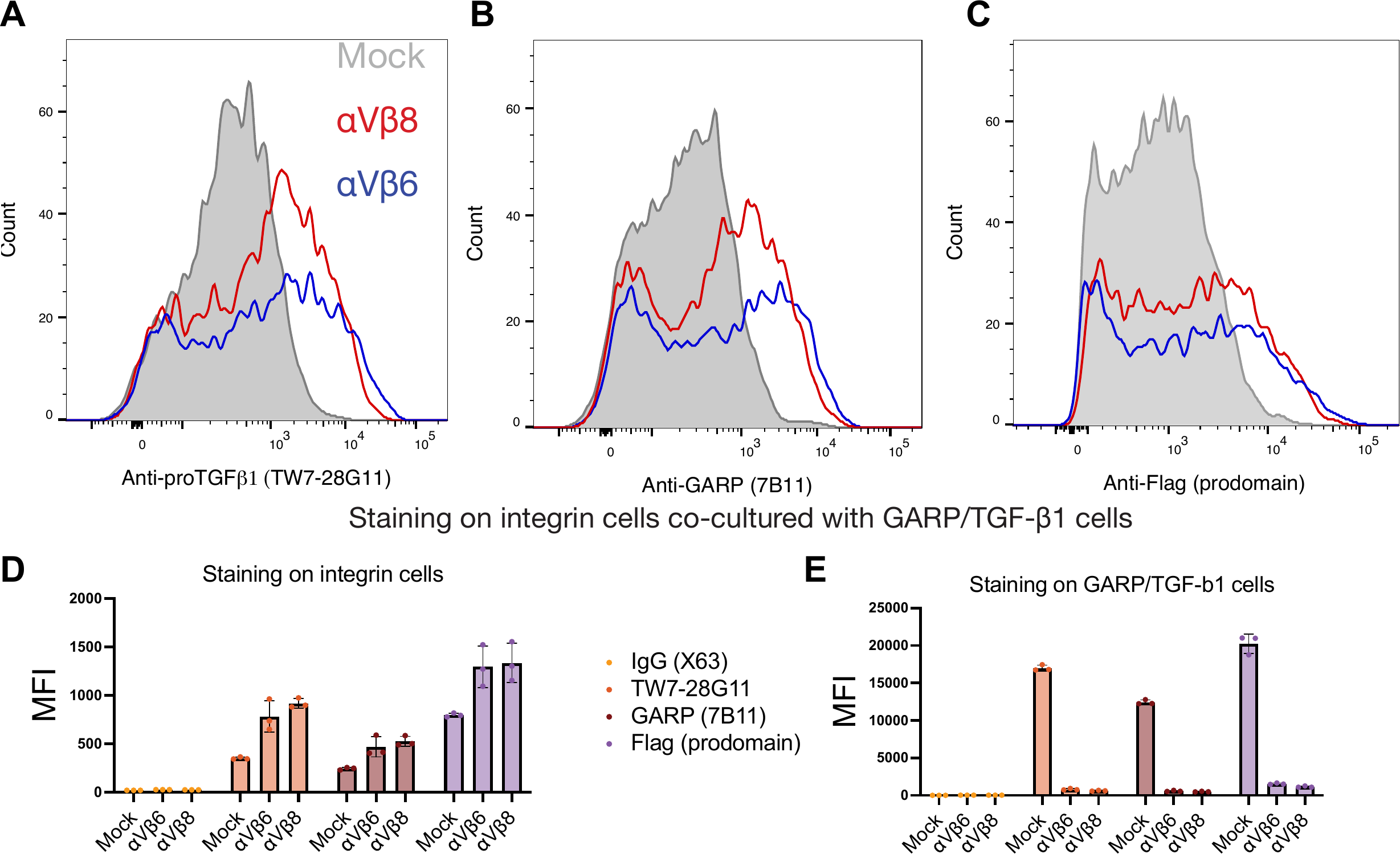
Intact GARP/TGF-β1 complexes are transferred to the surface of cells expressing activating integrins. A-C. Representative flow cytometry histograms of mock and integrin transfectants co-cultured with GARP/TGF-β1 transfectants and stained with the TW7-28G11 antibody specific for proTGF-β1 (A), an anti-GARP antibody (B), or an anti-Flag antibody (C), showing surface staining on integrin-expressing cells. D. Quantification of median fluorescence intensity of indicated antibodies on indicated integrin transfectants co-cultured with GARP/TGF-β1 transfectants showing surface accumulation of proTGF-β1 and GARP. E. Quantification of surface staining by indicated antibodies on GARP/TGF-β1 transfectants co-cultured with indicated integrin transfectants showing surface depletion of proTGF-β1 and GARP.

Similarly, we stained non-permeabilized cells with an anti-GARP antibody since the JF657::HaloTag labelling system to track Halo-GARP cannot distinguish between internalized from surface material on integrin-expressing cells by flow cytometry. In addition, since JF657 is cell-permeable^35^, it is expected to label both mature GARP complexed to TGF-β1 on the cell surface as well as immature forms at various stages of processing in the endomembrane system. GARP antibody staining showed the presence of GARP on the surfaces of integrin-expressing cells (Fig. 3B, D), and the loss of surface GARP from GARP/TGF-β1 transfectants co-cultured with integrin-expressing cells (Fig. 3E). Both TW7-28G11 and the GARP antibody staining profiles are consistent with those from staining the prodomain with an anti-Flag antibody (Fig. 3C-E). Taken together, these results indicate that GARP/TGF-β1 complexes are transferred, apparently intact, from the surface of GARP/TGF-β1 expressing cell onto the surface of cells expressing activating integrins.

### Integrin cytodomains are dispensable for TGF-β1 uptake

Activation of latent TGF-β1 by αVβ6 is thought to occur through transmission of cytoskeletal force that unfolds the prodomain and releases the GF ^37^. The cytoplasmic tail of the β6 integrin subunit has NXXY motifs that serve as binding sites for the cytoplasmic adaptors talin and kindlin, and mutation of these motifs reduces the ability of integrin αVβ6 to activate TGF-β1 ^11,24^. The role of cytoplasmic adaptors in activation by integrin αVβ8 is less clear. The cytoplasmic tail of the β8 subunit is highly divergent from that of β6 and lacks motifs resembling the talin/kindlin binding sites (Supplementary fig. 3A). However, a distinct set of adaptors has been implicated in integrin αVβ8 function in studies of mice ^40–42^. At the same time, cell culture work suggests that the cytoplasmic tail of integrin αVβ8 is dispensable for TGF-β activation in some contexts ^43^.

To examine the role of the β6 cytoplasmic adaptors, we used a β6 construct with Tyr-to-Ala mutations in both NXXY talin/kindlin binding motifs (YYAA mutant), and a β6 construct truncated at residue 758, which removes both motifs (Δ758). αVβ6 YYAA and αVβ6 Δ758 showed similar expression levels to WT αVβ6 by αV-integrin antibody staining in flow cytometry. We truncated the cytoplasmic tail of the β8 subunit at position 710, retaining only the four most membrane-proximal residues. The truncation greatly reduced expression, with αVβ8 Δ710 being expressed at ∼15% of WT levels.

αVβ8 Δ710 and the αVβ6 mutants mediated the uptake of Halo-GARP and Flag-prodomain from GARP/TGF-β1 transfectants at levels comparable to the full-length integrins (Fig. 4A,B) and depleted Halo-GARP and prodomain from GARP/TGF-β1 transfectants relative to mock transfectants (Fig. 4C,D). GARP/TGF-β1 cells retained more GARP and prodomain remaining on their surfaces when co-cultured with αVβ8 Δ710 vs full-length αVβ8 transfectants, which may reflect greatly reduced expression of Δ710. These results indicate that the cytoplasmic domains of β6 and β8, and by extension their cytoplasmic adaptor interactions, are not required for the uptake of GARP/TGF-β1 complexes by integrin-expressing cells.

**Figure 4.**
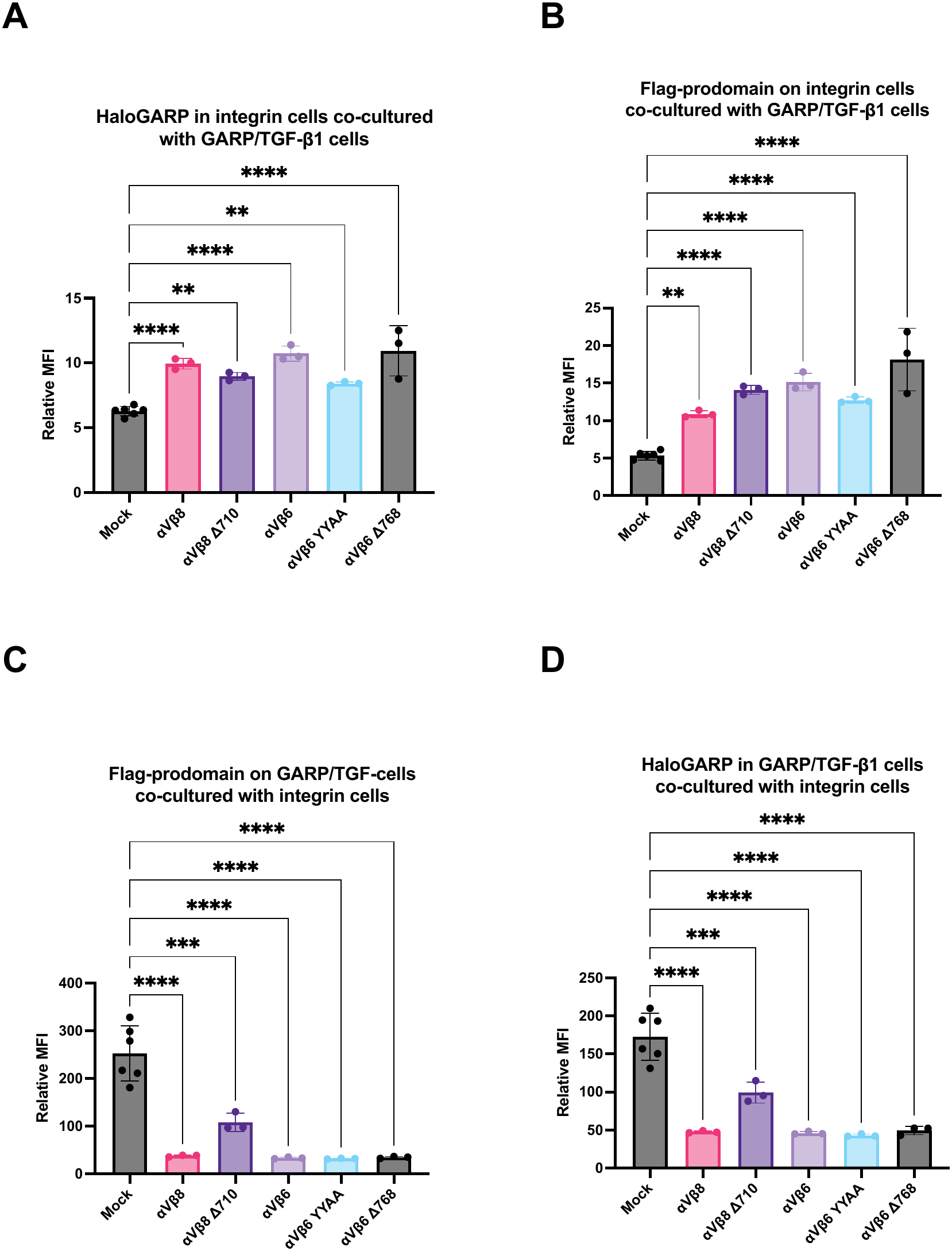
Integrin cytoplasmic tails are dispensable for trogoctotic transfer of GARP/TGF-β1 complexes. A, B. Quantification of JF657::Halo-GARP (A) and Flag-prodomain (B) signals on cells transfected with indicated integrins and co-cultured with Halo-GARP/TGF-β1 transfectants. C, D. Quantification of JF657::Halo-GARP (A) and Flag-prodomain (B) signals on Halo-GARP/TGF-β1 transfectants co-cultured with indicated integrin transfectants.

### Bystander cells are more sensitive to TGF-β blockade than TGF-β1 producing and activating cells

We next asked whether the transfer of latent TGF-β complexes to integrin-bearing cells has consequences for TGF-β signaling, and whether signaling differs between interacting TGF-β-producing cells and integrin-expressing cells and “bystander” cells that express neither TGF-β nor integrins. Traditional assays of TGF-β activation primarily rely primarily on reporter measurements in bystander cells. We therefore developed a tripartite reporter assay in which each activation condition is sequentially assayed in three configurations: with the reporter in GARP/ TGF-β1 cells, in integrin-expressing cells and in bystander cells. This design reveals cell type-specific differences in activation. Using this assay, we tested whether the three cell types responded equally to antibodies that neutralize the mature TGF-β GF: 1D11^44^ and Fresolimumab (GC1008) ^45^.

1D11 inhibited signaling similarly in TGF-β1-expressing and integrin-expressing cells for both αVβ6- and αVβ8-mediated activation (Fig. 5A,B). However, bystander cells were more sensitive to 1D11, with a 3-6-fold lower IC50 than TGF-β and integrin-expressing cells (Fig. 5G). Inhibition by Fresolimumab showed a similar pattern, with a 3-12-fold lower IC50 in bystander cells than in the interacting cell types (Fig. 5C,D,G). Whereas TGF-β1-producing and integrin-expressing cells interact physically, resulting in transfer of TGF-β1 complexes, bystander cells must receive active GF by diffusion from the site of activation. This requirement for diffusion may make bystander cells more sensitive to agents that neutralize soluble TGF-β1 GF.

**Figure 5.**
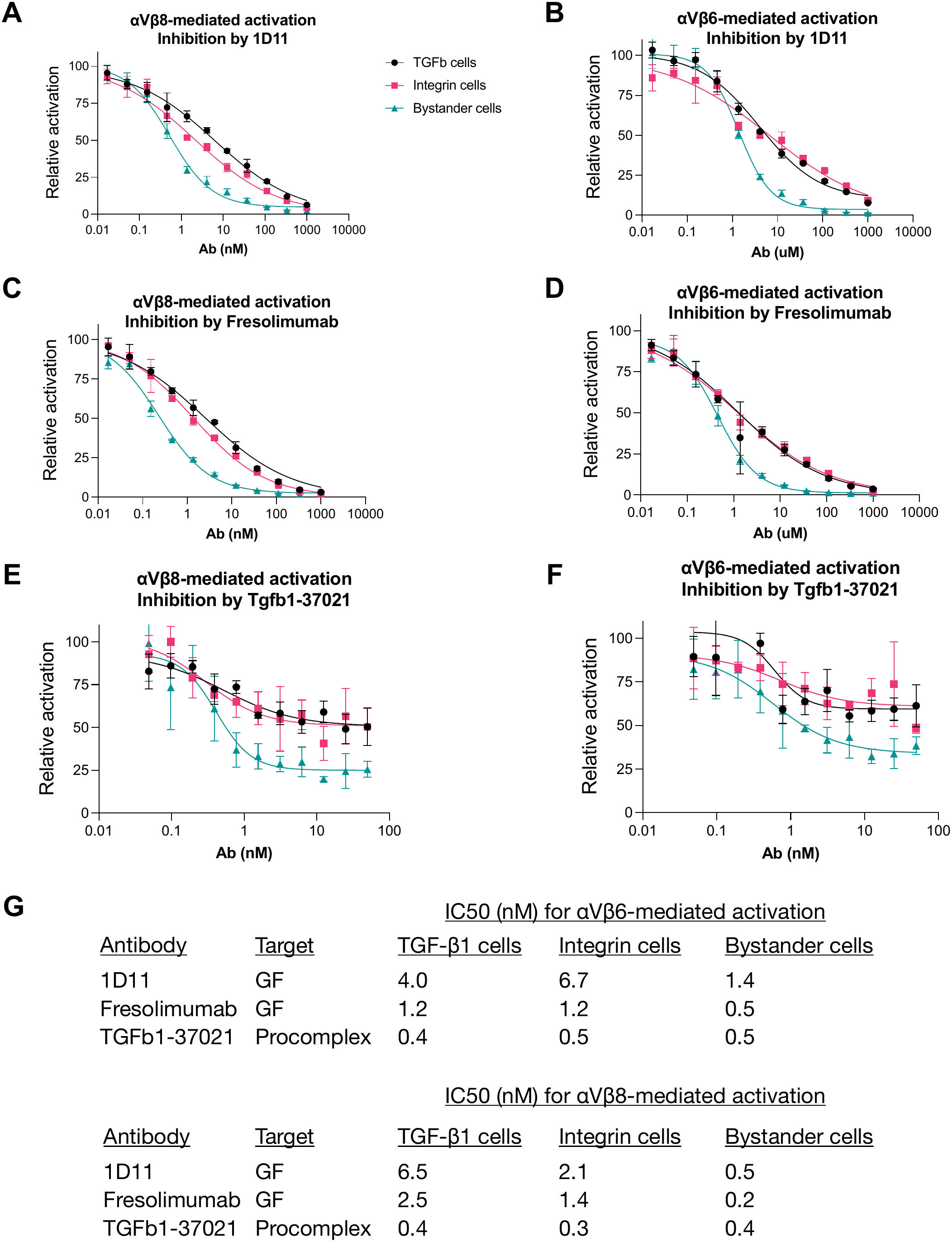
Bystander cells, which do not make or trogocytose TGF-β1, are more sensitive to GF-neutralizing antibodies than TGF-β1-expressing or integrin-expressing cells. A. CAGA-reporter assay showing inhibition of αVβ8-mediated GARP/TGF-β1 activation in TGF-β1-expressing cells (black), integrin-expressing cells (pink), and bystanders (green) by GF-targeting antibody 1D11. B. CAGA reporter assay showing stronger inhibition by 1D11 of αVβ6-mediated GARP/TGF-β1 activation in bystander cells. C, D. CAGA reporter assay showing stronger inhibition by Fresolimumab of αVβ8-mediated (C) and αVβ6-mediated (D) GARP/TGF-β1 activation in bystander cells. E, F. CAGA reporter assay showing inhibition by latent TGF-β1-neutralising antibody Tgfb1-37021 of αVβ8-mediated (E) and αVβ6-mediated (F) GARP/TGF-β1 activation in different cell types. G. Table of IC50 values for GF-targeting and latent complex-targeting antibodies in TGF-β1-expressing cells, integrin-expressing cells, and bystander cells.

To test this hypothesis further, we measured the effect of Tgfb1-37021, an antibody that binds latent TGF-β1 and prevents its activation ^46^. Since this antibody blocks activation prior to the GF diffusion step, we predicted that it would affect all cell types equally. Consistent with our hypothesis and in contrast with 1D11 and Fresolimumab, Tgfb1-37021 inhibited signaling in all three cell types with similar IC50s (Fig. 5G). Intriguingly, the extent of inhibition was about two-fold greater in bystander cells than in TGF-β1-expressing and integrin-expressing cells (Fig. 5E,F). This could reflect partial shielding of latent complexes from extracellular antibodies on trogocytosing cells, which remains to be tested.

### TGF-β1 uptake can activate the downstream TGF-β pathway

We next examined mutations that block furin cleavage, which is expected to prevent GF release. The furin recognition sequence RHRR in TGF-β1 was mutated to RHRA (R249A) or AHAA. Neither the R249A nor the AHAA mutation affected transfer of GARP/TGF-β1 complexes from TGF-β1-expressing cells to integrin-expressing cells (Fig. 6A-H). We then tested whether the mutants could activate downstream signaling in TGF-β1-expressing and integrin-expressing cells, compared with bystander cells. Both mutants produced significant activation in TGF-β1-expressing cells (Fig. 6I) and integrin-expressing cells (Fig. 6J), but not in bystander cells (Fig. 6K), although the magnitude of the response was lower than with WT. Mutations in the furin cleavage site therefore do not attenuate GARP/TGF-β1 trogocytosis. They abolish signaling in bystander cells while attenuating signaling in TGF-β1 and integrin-expressing cells. Lack of cleavage is expected to keep the GF tethered to the prodomain, restricting signaling to TGF-β receptors on the same cell. Signaling can therefore occur on the TGF-β1-expressing cell and on integrin-expressing cells that have taken up TGF-β1 complexes, whereas the inability of the GF to diffuse precludes signaling to bystander cells.

**Figure 6.**
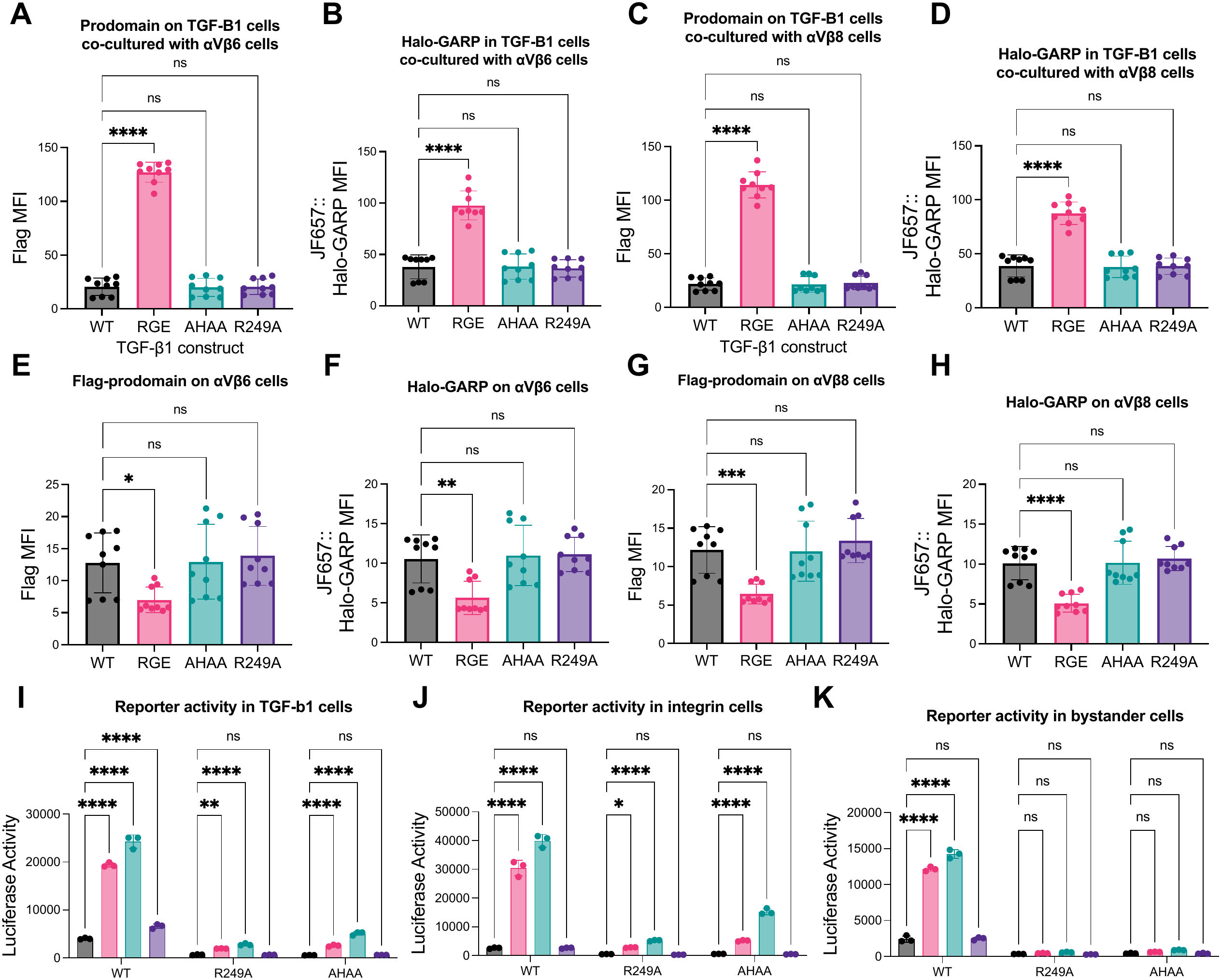
Furin cleavage site mutations do not attenuate GARP/TGF-β1 trogocytosis, attenuate but do not abolish signaling in TGF-β1 and integrin-expressing cells, and abolish signaling in bystander cells. A, B. Quantification of Flag-prodomain (A) and Halo-GARP (B) signal in Expi293F cells transfected with GARP and the indicated TGF-β1 constructs and co-cultured with integrin αVβ6 transfectants showing that furin cleavage mutants are depleted similarly to WT TGF-β1. C, D. Quantification of Flag-prodomain (C) and Halo-GARP (D) signal in Expi293F cells transfected with integrin αVβ6 co-cultured with cells transfected with GARP and the indicated TGF-β1 constructs showing that furin cleavage mutants are trogocytosed similarly to WT TGF-β1. E, F, G, H. Quantification of Flag-prodomain (E,G) and Halo-GARP (F,H) in integrin αVβ8 transfectants and GARP/TGF-β1-expressing cells as in A-D. I. Reporter assay showing activation of CAGA-luciferase reporter activity in GARP/TGF-β1 transfectants expressing indicated TGF-β1 mutants co-cultured with cells expressing indicated integrins. J. Reporter assay showing activation of CAGA-luciferase reporter activity in integrin-expressing cells co-cultured with GARP/TGF-β1 transfectants expressing indicated TGF-β1 mutants. K. Reporter assay showing activation of CAGA-luciferase reporter activity in bystander cells co-cultured with integrin-expressing cells and GARP/TGF-β1 transfectants expressing TGF-β1 mutants.

### Conversion of integrin-expressing cells into TGF-β signaling relays

Does the pool of latent GARP/TGF-β1 complexes taken up by integrin-expressing cells participate in signaling? To test this, we pre-incubated CMFDA-labeled mock or integrin transfectants with GARP/TGF-β1 transfectants for 90 minutes, sorted the CMFDA+ population of integrin cells, and tested their ability to signal to reporter cells (Fig. 7A, Supplementary fig. 4A). Reporter cells co-cultured with mock transfectants showed no signaling under any condition. Reporter cells co-cultured with integrin αVβ6 or αVβ8 transfectants showed robust activation only when the transfectants were pre-incubated with GARP/TGF-β1 cells (Fig. 7B). This suggested that the integrin-expressing cells could take up GARP/TGF-β1 complexes and subsequently activate and transmit the GF to other cells, turning them into TGF-β signaling relays. To exclude reporter activity arising from contaminating GARP/TGF-β1-expressing cells in the sorted population, we determined the fraction of contaminating GARP/TGF-β1-expressing cells that would be needed in a population of integrin αVβ8 transfectants to produce activity similar to that of sorted αVβ8 transfectants pre-incubated with GARP/TGF-β1-expressing cells. About 6% contamination would be required, compared with the measured contamination of ∼0.5% (Supplementary fig. 4C-E), confirming that the reporter activity reflects a bona fide contribution from integrin-expressing cells.

**Figure 7.**
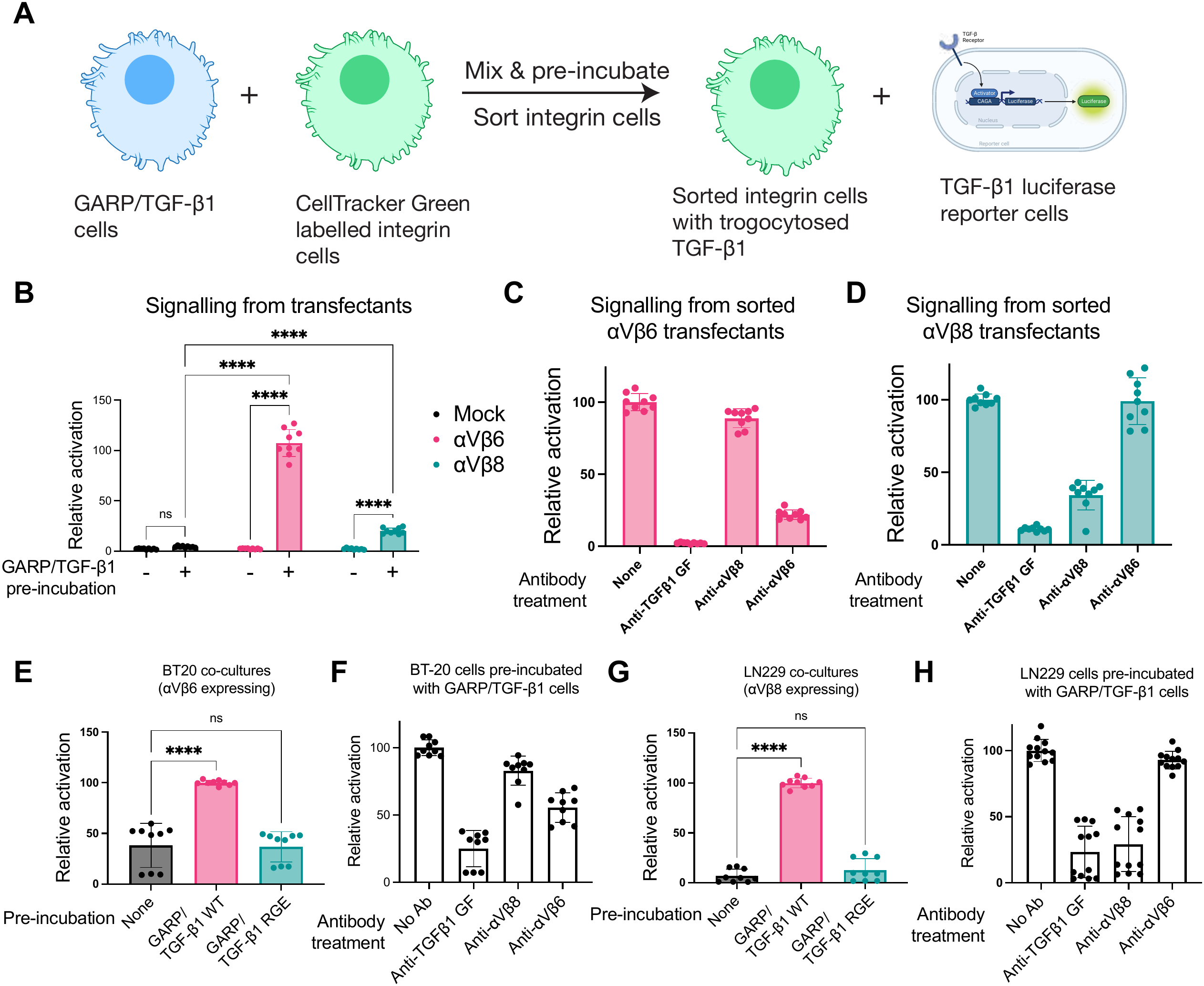
Trogocytosis converts integrin-expressing cells into TGF-β1 signaling relays. A. Schematic of assay to study signaling from trogocytosed material on integrin-expressing cells. Integrin cells are pre-incubated with GARP/TGF-β1 transfectants, sorted, and then co-cultured with reporter cells. B. CAGA-luciferase reporter activity in co-cultures of reporter cells with mock or integrin Expi293F transfectants sorted with or without pre-incubation with GARP/TGF-β1 transfectants. Integrin transfectants robustly induced reporter activity, but only when pre-incubated with GARP/TGF-β1 cells. C, D. CAGA reporter assay showing effect of GF and integrin-blocking antibodies on signaling from αVβ6 (C) or αVβ8 (D) cells pre-incubated with GARP/TGF-β1 cells. E. CAGA-luciferase activity from co-cultures of reporter cells and αVβ6-expressing BT-20 cells sorted without pre-incubation, or pre-incubated with GARP/TGF-β1 WT or RGE transfectants. F. Reporter assay showing inhibition by integrin-blocking antibodies of signaling from BT-20 cells pre-incubated with GARP/TGF-β1 transfectants. G. CAGA-luciferase activity from co-cultures of reporter cells and αVβ8-expressing LN229 cells sorted without pre-incubation, or pre-incubated with GARP/TGF-β1 WT or RGE transfectants. H. Reporter assay showing inhibition by integrin-blocking antibodies of signaling from LN229 cells pre-incubated with GARP/TGF-β1 transfectants.

We further characterized this signaling relay using different inhibitory antibodies. Signaling from integrin αVβ6 or αVβ8 transfectants was blocked by Fresolimumab, consistent with the release of TGF-β1 GF from the integrin-expressing cells (Fig. 7C,D). Antibodies that neutralize integrin αVβ6 and αVβ8 specifically inhibited signaling from cells expressing the cognate integrins (Fig. 7C,D). This shows that TGF-β1 activation requires integrin engagement even after latent complexes are transferred to the integrin-expressing cell. We next asked whether the amount of GARP/TGF-β1 complex on an integrin-expressing cell determines its ability to drive bystander cell signaling. To address this, we repeated the sorting experiment by pre-incubating JF657-labeled Halo-GARP/TGF-β1 cells with CellTracker Green-labeled αVβ6 or αVβ8 transfectants. After gating on the CellTracker channel, we used the JF657 channel to gate integrin cells into Halo-GARP high, medium and low populations (Supplementary Fig. 5). Signaling from both αVβ6 and αVβ8 transfectants was driven primarily by the JF657^high^ population (Supplementary Fig. 5), suggesting that the strength of signaling to bystander cells increases with the number of GARP/TGF-β1 complexes taken up by a cell.

Finally, we tested the ability of BT-20 and LN229 cells to act as TGF-β relays. Sort purity exceeded 99.4% (Supplementary Fig. 4F,G). For both cell lines, pre-incubation with GARP/TGF-β1-expressing cells produced significantly greater TGF-β activation than with no pre-incubation or pre-incubation with GARP/TGF-β1 RGE-expressing cells (Fig. 7E,G). Inhibition by Fresolimumab and anti-integrin antibodies confirmed GF release and that integrin binding was required (Fig. 7F,H).

## Discussion

In this study, we tracked the fate of all members of the GARP/TGF-β1 complex during activation by integrins αVβ6 and αVβ8. This led to the surprising discovery that TGF-β1 complexes are transferred intact to integrin-expressing cells during activation. This process was very quick and triggered by direct cognate receptor-ligand interactions between the prodomain and integrins. The transfer of GARP/TGF-β1 complexes to integrin cells bore several hallmarks of trogocytosis or “cell nibbling”, a process by which surface molecules are transferred from one cell to another cell by cell-cell contacts^47,48^. As in other reports of trogocytosis^49–51^, significant transfer of GARP/TGF-β1 complexes to integrin cells occurs in just 10 minutes, and requires direct receptor-ligand interactions between participating cells as shown by the lack of transfer to cells not expressing integrins or those expressing the RGE mutant TGF-β1. TGF-β1 trogocytosis is observed in Expi293F cells transiently transfected with integrins, and in cancer cells that natively express TGF-β1-activating integrins, suggesting it is a general feature of integrin-dependent activation.

Previous studies on TGF-β1 activation assumed that the GF was primarily trafficked by diffusion from the site of activation. Studies on TGF-β1 activation primarily relied upon the use of reporter cell lines that measure bulk TGF-β1 GF concentrations in the medium^17,18,24,52,53^. These methods ignore signaling in the cells that produce and activate TGF-β1 and instead only focus on signaling in cells that are bystanders to trogocytosis and TGF-β1 activation. To understand how trogocytosis affects downstream TGF-β1 pathway responses in TGF-β1 expressing cells, integrin expressing cells, and bystander cells, we extended the traditional bystander cell-based reporter assay into a tripartite assay that examines signaling in all relevant cell types. We find that different cell types have different levels of sensitivity to GF inhibition, with signaling in bystander cells inhibited at lower doses than in integrin or TGF-β1-expressing cells. Our results suggest that the dosing of these antibodies may need to be adjusted not just based on how much inhibition is desired, but also based on which cell types need to be therapeutically targeted.

TGF-β1 receptors are expressed ubiquitously, yet the effects of TGF-β1 signaling tend to be extremely cell-type specific ^54^. Genetic evidence from mice suggests that TGF-β1 is presented in different tissues in complex with specific “milieu anchors” that are thought to provide highly selective and localized activation of TGF-β1^17^. Using our tripartite reporter assay, we found that furin/proconvertase cleavage was essential for activation in bystander cells, but not in integrin or TGF-β1-expressing cells. This suggests that trogocytosis could be an important mechanism of TGF-β1 trafficking in milieus where activating integrins are expressed, ensuring downstream signals are localized to cells that can activate TGF-β1. Our findings are consistent with a recent report that mice bearing an equivalent of the R249A mutation in TGF-β1 do not show the lethal inflammatory phenotypes reported in TGF-β1 null mice^43^.

How is latent TGF-β1 presented on cell surfaces activated by integrins? We found that integrin-neutralizing antibodies could block the transfer of latent TGF-β1 complexes to cells expressing activating integrins and prevent activation and signaling. However, these antibodies could also block signaling from integrin-expressing cells that have already taken up latent TGF-β1 complexes. These results point to an initial and rapid integrin-mediated transfer of latent TGF-β1 complexes to the integrin-expressing cell, followed by a slower process in which the complexes are activated on the integrin-bearing cell. The processing in the integrin-bearing cell results in the activation of downstream signaling not just in the integrin-bearing cell, but also in nearby bystander cells via diffusion of the GF. We find that mutations that block prodomain-GF cleavage, and therefore diffusion, block signaling in bystander cells but not in integrin or TGF-β1 bearing cells. How can signaling be activated in the trogocytosing cells? Recent structural work on TGF-β1-αVβ8 complexes was used to propose a model of binding-induced conformational changes in latent TGF-β1 being sufficient to induce enough flexibility in the structure to allow the GF to bind receptor complexes without dissociating from the prodomain^55^. However, subsequent structural analysis showed that the TGF-β1-αVβ8 complexes lacked the conformational changes required in the α3 helix for interactions with TGFβR1 due to clashes with the α1 helix^56^. Cellular forces transmitted through the cytoplasmic tail adaptors of the β6 subunit can cause deformations in the α1 helix to allow the GF to access the receptor^37^, but this model does not explain the ability of integrins αVβ6 and αVβ8 to activate TGF-β1 without their β-subunit cytodomains. One possibility is that the trogocytosed TGF-β1 is subsequently processed by integrin-mediated internalization, following which dissociation of the prodomain and GF may happen following proteolytic processing or through interactions with other proteins in an internal cellular compartment. While it has not been studied in the context of αVβ8, integrin αVβ6 can drive the internalization of ligands including soluble latent TGF-β1 complexes^57^. Studies using αVβ6-targeting antibodies suggest that internalization is mediated by integrin binding to an RGD motif ^58^. Studies on TGF-β receptor function have shown that they are primarily activated in endosomal compartments ^59–61^. Trogocytosis of latent TGF-β complexes may thus facilitate their co-localization with receptors in the integrin-expressing cell.

In conclusion, to our knowledge our study provides the first report of the transfer of latent TGF-β complexes to integrin cells. Our findings establish this transfer as an important step in activation and show that it is capable of converting integrin-expressing cells into “signalling relays” that transmit TGF-β signals even in the absence of the original TGF-β producing cell. With our new tripartite reporter assay, we identify important differences in responses of different cell types to TGF-β activation and to TGF-β inhibitors.

## Materials and methods

### Cell culture

Expi293F cells (Gibco, A14527) were cultured in suspension in Expi293 expression medium (Gibco, A1435101). For transfection they were plated in FreeStyle 293 Medium (Gibco, 12338018). Expi293F cultures were grown at 37℃ in 8% CO2. CAGA reporter cells^52^ were a gift from Tom Thompson (University of Cincinnati) and were cultured in DMEM (Gibco, 11995065) supplemented with 10% fetal bovine serum (FBS) and 100 μg/ml G418 37℃ in 7.5% CO_2_. LN-229 cells were purchased from ATCC (CRL-2611) and were cultured in DMEM with 10% FBS at 37℃ in 5% CO_2_. BT-20 cells were a gift from Judy Lieberman (Boston Children’s Hospital) and were cultured in EMEM (ATCC, 30-2003) with 10% FBS at 37℃ in 5% CO_2_. CAGA-firefly reporter cells were constructed by replacing the dynGFP sequence in CAGA-dynGFP{Marvin, 2022, 39521} with firefly luciferase. Stable Expi293F reporter lines were established with puromycin selection. Cells were then clonally isolated and their sensitivity to TGF-β1 GF determined by a GF standard curve. The clone with the highest dynamic range was picked for reporter assays.

### Antibodies

Integrin staining and function blocking antibodies used were 17E6 (anti-αV)^62^, 7.1G10 (anti-β6)^58^, ADWA11 (anti-β8)^63^. For immunostaining they were used with a Goat anti-mouse AlexaFluor 647 secondary antibody (Invitrogen, A-21237). 1D11 was purified from hybridoma (ATCC, HB-9849) supernatant. Sequences of Tgfb1-37021^46,64^ and Fresolimumab^65^ were identified from their respective patents and converted to DNA sequences using IDT’s codon optimization tool. They were cloned into pD2529-based expression vectors. They were expressed via transient transfection in Expi293F cells and purified from supernatants. GARP immunostaining was done using anti-human GARP antibody clone 7B11 (Biolegend, 352502). Flag staining was done using Brilliant Violet 421 anti-DYKDDDDK Tag Antibody (Biolegend, 637322).

### Expression constructs

Expression constructs for GARP, Flag-TGF-β1, integrin αV subunit, integrin β6 and β8 subunits have been described previously^24^. Mutations or truncations were generated by Gibson assembly with mutant sequences ordered from IDT as gBlocks. For the Halo-GARP construct, the HaloTag7 sequence^33^ was inserted immediately after the signal peptide sequence. K344*, RGE, R249A, and AHAA mutations were made by amplifying the region around the mutation with PCR primers containing the desired mutation and flanking sequences, followed by Gibson assembly.

### Fluorescent labelling

For site-specific labelling of the GF, cells were transfected with plasmids encoding Halo-GARP and Flag-TGF-β1 with an amber codon at position K344 (Flag-TGF-β1 K344*), along with pCMV_NESPylRS(AF)_hU6tRNAPyl (Addgene #223508)^25^, which encodes a pyrrolysine tRNA and a mutated tRNA synthase that accepts the TCO*-lys residue, and pcDNA5-eRF1-E55D, which encodes a eukaryotic release factor 1 mutant that decreases polypeptide termination^27^. After transfection, cells were incubated with 25μM TCO*-lys (SiChem GmbH, SC-8008) for 48h. Before labelling, cells were incubated in fresh FreeStyle 293 media to remove excess TCO*-lys. Cells were incubated with 500 nM LD655-tetrazine (Lumidyne) for 1 hour under standard culture conditions, washed 3x in PBS, and resuspended in FreeStyle 293 media. Labelling of mock transfectants not expressing any amber mutant construct was used to determine the background.

Halo-GARP labelling was done with JF657^35^. JF657 was a gift from Luke Lavis (HHMI/Janelia). 1 nmol of JF657 was resuspended in 10 μL of anhydrous DMSO and diluted into FreeStyle 293 media to a final concentration of 200 nM for labelling. Halo-GARP transfected cells were incubated with JF657 for 1 hour under standard culture conditions, washed 3x in PBS, and resuspended in FreeStyle 293 media. Labelling of mock transfectants not expressing Halotag was used to ensure specificity of labelling conditions.

Cells were stained with 10 μM CellTracker Green CMFDA (Invitrogen, C7025) for 15 min in FreeStyle 293 media. They were then washed thrice in PBS and resuspended in FreeStyle 293 media.

### Immunostaining and Flow cytometry

After co-culture, cells were resuspended in ice-cold staining buffer (HBSS, supplemented with 1% BSA, 1mM CaCl_2_, 1mM MgCl_2_) and stored and stained on ice until readings were made by flow cytometry. All unconjugated primary antibodies were diluted in staining buffer to a final concentration of 2 μg/ml for flow cytometry. Cells were incubated in primary antibody for 1 hour, washed 3x in staining buffer, incubated with secondary antibody (diluted 1:500) for 1 hour, washed 3x in staining buffer, and resuspended in staining buffer prior to measurement. Flow cytometry was done on BD FACSCanto II Flow cytometer. Analysis was done using FlowJo v10.10.2.

### Cell sorting-based signalling experiments

Mock or integrin-expressing cells were stained and resuspended in FreeStyle 293 medium at a concentration of 10^6^ cells/ml. GARP/TGF-β1 cells were resuspended without labelling in FreeStyle 293 medium at a concentration of 10^6^ cells/ml. Equal volumes of both cell types were mixed and pre-incubated for 1.5 hours at 37℃. Samples were then kept on ice until sorting was complete. Sorting was performed on a Sony SH-800Z cell sorter. Data was collected and analysed using Sony Cell Sorter Software v2.1.7. A 100 μm sorting chip was used for sorts involving Expi293F transfectants, and a 130 μm chip was used to sort LN-229 and BT-20 cells. Cells were sorted into FreeStyle 293 media. After counting, sorted cells were seeded at a density of 15,000 cells/well into wells containing 15,000 CAGA cells each. Luciferase readings were done 24h after co-culture using the Promega luciferase assay system (Promega, E1500).

## Acknowledgements

We thank Prof. Peter ten Dijke (Universiteit Leiden) for the CAGA-dynGFP reporter and Prof. Judy Lieberman (Boston Children’s Hospital) for BT-20 cells. This work was supported by NIH Grant HL159714 (T.A.S.), NIH Grant DK124443 (V.Q.L.), and Cancer Research Institute Irvington Postdoctoral Fellowship CRI5057 (A.B.R).

## Notes

### Competing Interest Statement

The authors have declared no competing interest.

