## Supplementary Figures for "Intercellular trafficking of latent TGF-β1 converts integrin-expressing cells into signaling relays"

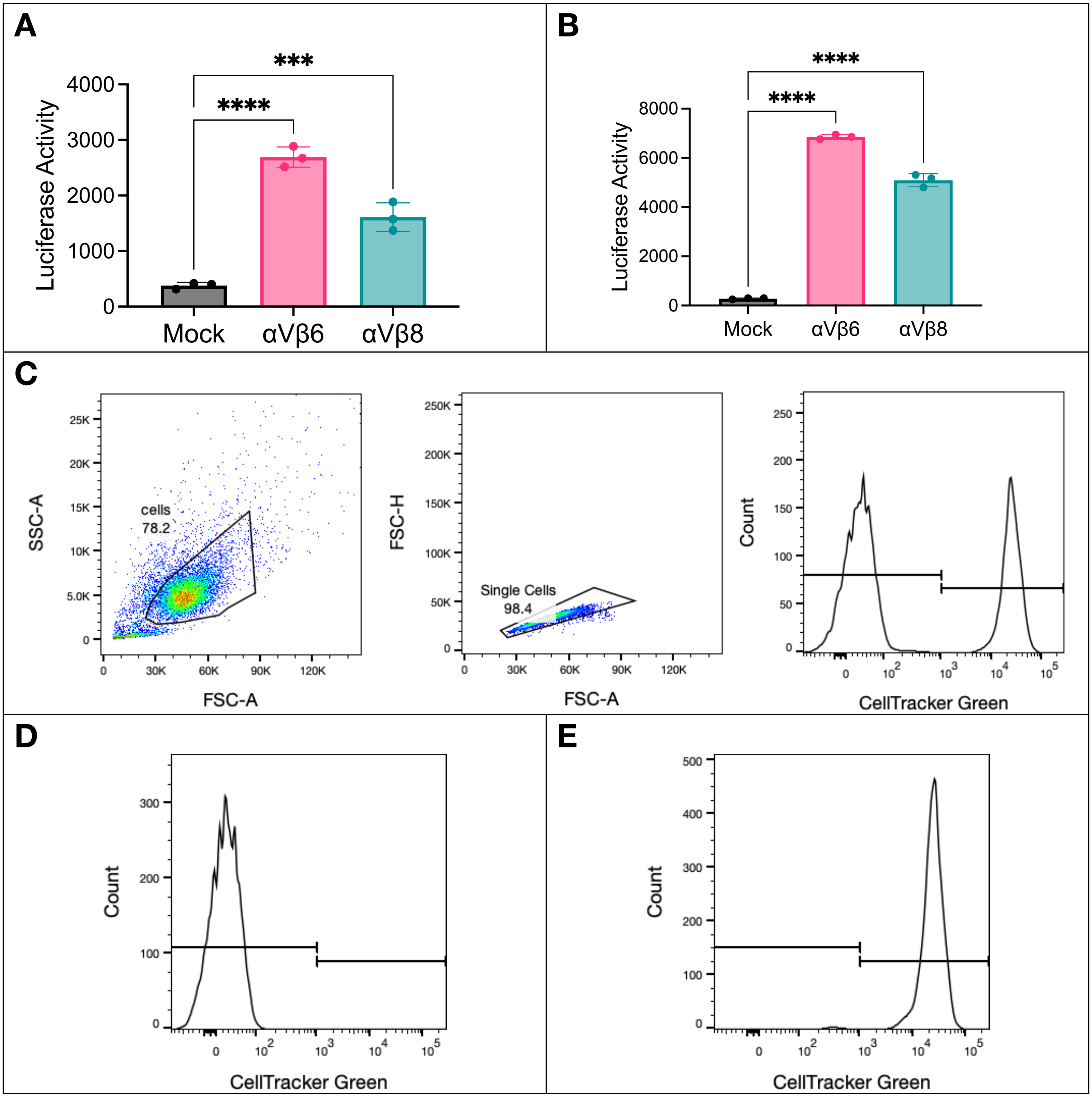


**Supplementary Figure 1: Controls for GARP/TGF-β1 tracking experiments.**

A. Activation assay with LD655-labelled GARP/TGF-β1 K344* transfectants co-cultured with mock or integrin transfectants.

B. Activation assay with JF657-labelled HaloGARP/TGF-β1 transfectants co-cultured with mock or integrin transfectants.

C. Sample gating strategy for the analysis of GARP/TGF-β1 trafficking to integrin-expressing cells. Cells are gated on forward and side-scatter to eliminate debris, then gated between forward scatter area and height to select for single cells. Finally CellTracker Green signal is used to identify GARP/TGF-β1 and integrin cells.

D, E. Controls without co-culture for CellTracker Green staining. Unstained GARP/TGF-β1 cells (D) are negative, while stained integrin cells are positive (E).

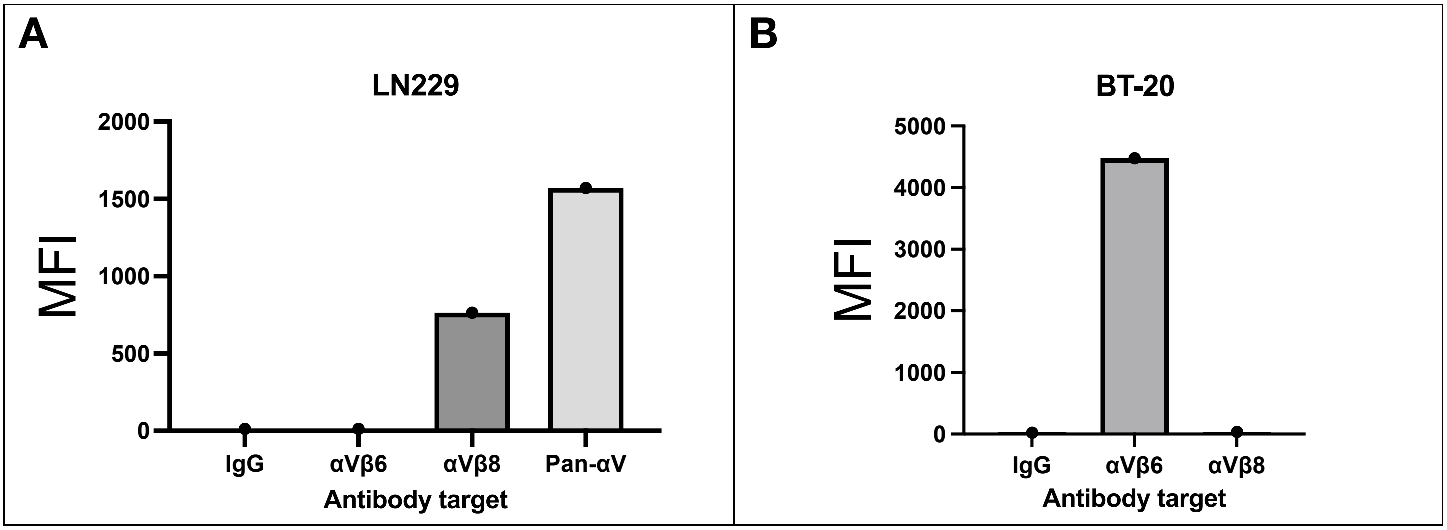


**Supplementary Figure 2: Expression of integrins on cancer cell lines.**
Immunostaining of LN229 (A) and BT-20 (B) cell lines with antibodies targeting αVβ6 and αVβ8, showing αVβ8 expression on LN229 cells and αVβ6 expression on BT-20 cells.

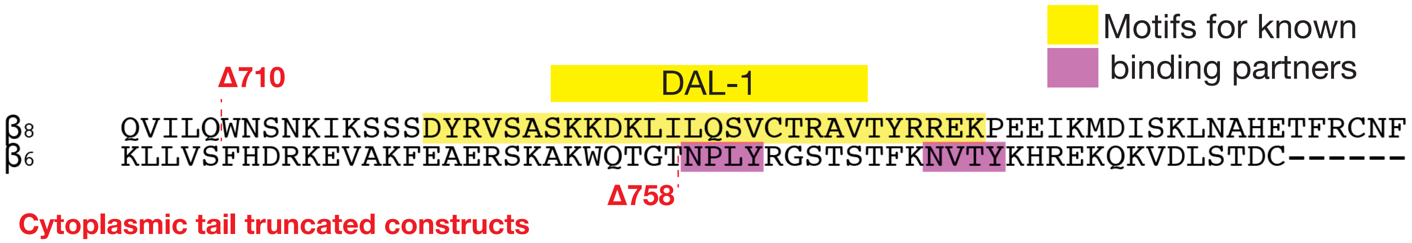


**Supplementary figure 3. Cytoplasmic tail mutations in β6 and β8 subunits.**
Positions of truncations in β6 and β8 subunits are shown in the context of binding motifs for previously identified interactors.

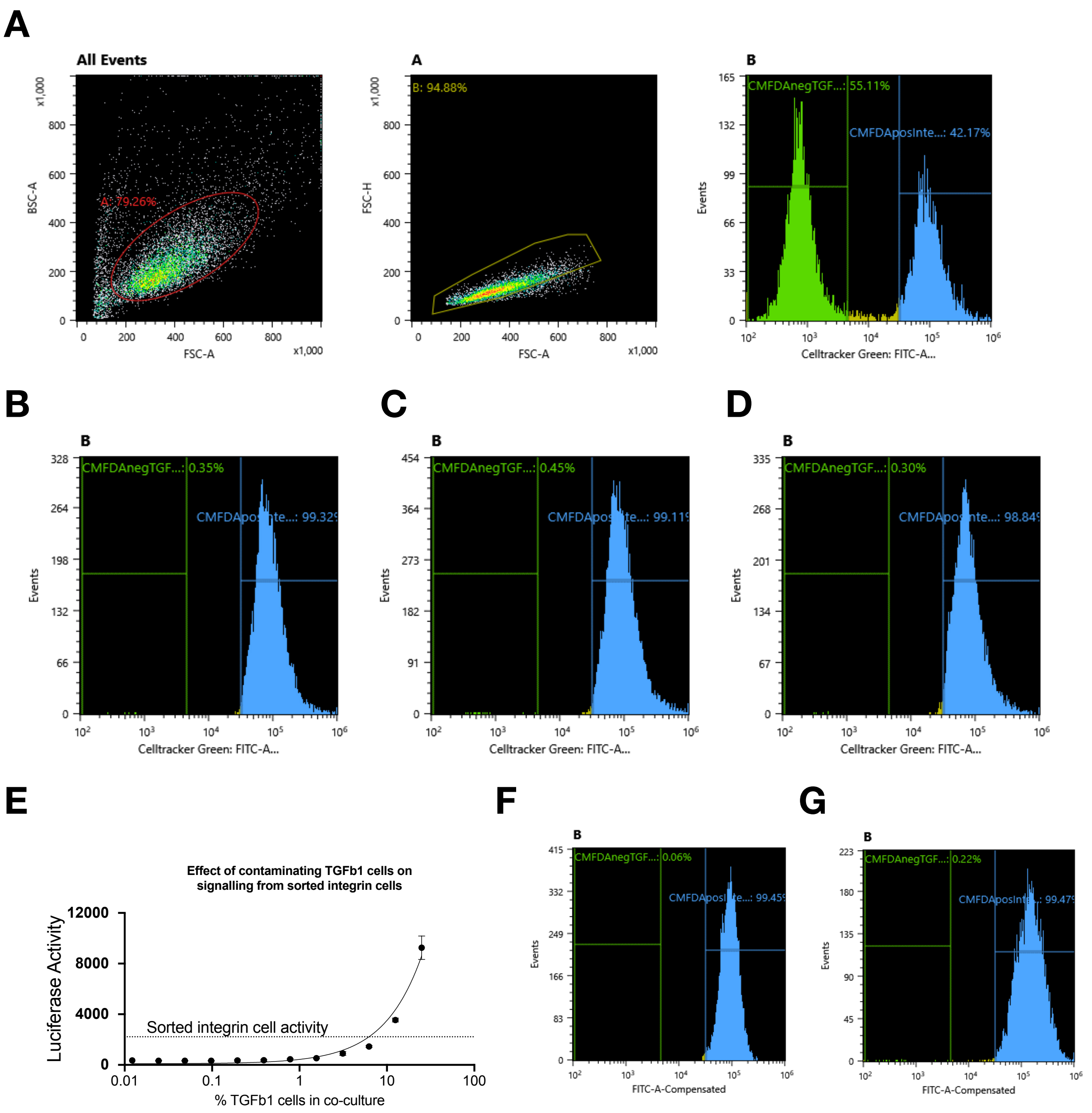


### Supplementary figure 4: Controls for sorting experiments

A. Sample gating strategy used to sort integrin-expressing cells after co-culture. After gating out debris and gating for singlets, CellTracker Green CMFDA staining is used to sort integrin cells.
B. Purity testing of sorted cells. Mock transfectants were co-cultured with GARP/TGF-β1 transfectants. Cells were sorted as described in (A). Sorted cells were re-run for analysis of purity, and the percentages of cells in the negative and positive gates were calculated as a measure of sort purity.
C. Purity testing of sorted integrin αVβ6 transfectants.
D. Purity testing of sorted integrin αVβ8 transfectants.
E. Contamination analysis. Integrin αVβ8 transfectants were mixed with GARP/TGF-β1 transfectants as indicated, and the level of signalling was measured with bystander cells. This was compared to the level of signalling from sorted integrin αVβ8 cells to test whether the activation of bystander cells could be explained by contaminating GARP/TGF-β1 cells.
F. Purity testing of sorted LN229 cells.
G. Purity testing of sorted BT-20 cells.

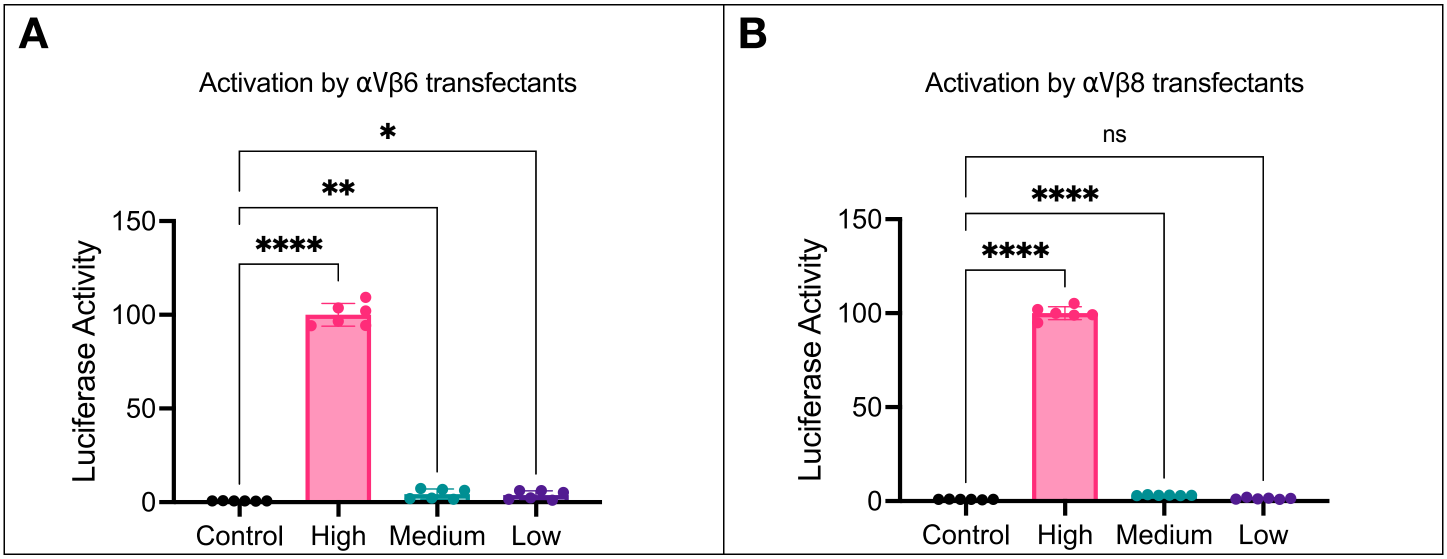


**Supplementary figure 5: Dose-dependence-relationship between trogocytosed material and ability to signal to bystander cells.**
CellTracker Green CMFDA stained αVβ6 (A) or αVβ8 (B) transfectants were pre-incubated with JF657-labelled HaloGARP/TGF-β1 transfectants. Integrin cells were sorted on the basis of CMFDA positivity and gated as high/medium/low based on JF657::HaloGARP staining. After sorting JF657 high/medium/low integrin cells were co-cultured with reporter cells along with mock sorted cells.
